# SWAMPy: Simulating SARS-CoV-2 Wastewater Amplicon Metagenomes with Python

**DOI:** 10.1101/2022.12.10.519890

**Authors:** William Boulton, Fatma Rabia Fidan, Hubert Denise, Nicola De Maio, Nick Goldman

## Abstract

**Motivation:** Tracking SARS-CoV-2 variants through genomic sequencing has been an important part of the global response to the pandemic. As well as whole-genome sequencing of clinical samples, this surveillance effort has been aided by amplicon sequencing of wastewater samples, which proved effective in real case studies. Because of its relevance to public healthcare decisions, testing and benchmarking wastewater sequencing analysis methods is also crucial, which necessitates a simulator. Although metagenomic simulators exist, none are fit for the purpose of simulating the metagenomes produced through amplicon sequencing of wastewater.

**Results:** Our new simulation tool, SWAMPy (**S**imulating SARS-CoV-2 **W**astewater **A**mplicon **M**etagenomes with **Py**thon), is intended to provide realistic simulated SARS-CoV-2 wastewater sequencing datasets with which other programs that rely on this type of data can be evaluated and improved.

**Availability:** The code for this project is available at https://github.com/goldman-gp-ebi/SWAMPy It can be installed on any Unix-based operating system and is available under the GPL-v3 license.

## Introduction

Wastewater sequencing has proven useful in the genomic surveillance of SARS-CoV-2 and can provide a less-biased picture of the variants circulating in a population than clinical surveillance (1). Amplicon sequencing is the preferred method for this purpose since it is efficient in terms of cost, labour and time, and is well-suited for heavily contaminated samples — as may be found with biological samples collected for SARS-CoV-2 sequencing — thanks to its targeted nature (2). Such sequencing has typically been done via multiplex PCR using a pre-defined primer set with paired-end reads generated by an Illumina device (1).

A number of methods and software tools for wastewater SARS-CoV-2 sequencing data analysis are available such as SAM Refiner (3), COJAC (4), LCS (5) and Freyja (6). Evaluating the effectiveness of new methods on *in vivo* or *in vitro* samples is often difficult or impossible, for example because of the lack of availability of a wide range of real or synthetic samples and the costs of repeated experiments (7). However, simulated datasets can provide an efficient way of bench-marking new methods’ performances (8).

There is a specific set of features characteristic to data coming from wastewater amplicon sequencing. For example, it has been shown that there is a high variation in amplification across different amplicons of a given primer set, resulting in a variation in read depth across the genome (1, 2). Moreover, wastewater data is expected to represent a mixture of different SARS-CoV-2 variants since the biological matter in the sample comes from multiple people, and will carry RNA degradation signatures resulting from the environmental exposure of the viral RNAs in sewage as well as PCR, sequencing library layout-specific and sequencing device-specific errors (9–11).

Existing standard metagenomic simulators do not attempt to capture all the characteristics seen in the amplicon sequencing protocols used for SARS-CoV-2 such as the ARTIC community protocols (12; https://artic.network/ncov-2019; https://artic.network/resources/ncov/ncov-amplicon-v3.pdf) widely used to prepare samples for Illumina sequencing platforms. For example, InSilicoSeq (13) is intended to produce shotgun metagenomic sequences; and while Grinder (7) can simulate amplicon sequencing, it cannot be tuned to produce a bespoke amplicon distribution and does not produce realistic sequencing quality scores. The simulation tool ART (14) can also generate reads for amplicons, but only in equal proportions. Studies (5, 15–17) have demonstrated the need for a dedicated wastewater SARS-CoV-2 sequencing simulator. Each, however, performed its own simulations for its specific use case, with most simulators limited to simplified scenarios with uniform amplicon abundances and only read errors (e.g. 15, 16, 18), or recreating the read-depth variation of one specific real experiment (e.g. 5, 17). Both cases have drawbacks, either risking overfitting to one dataset, or omitting important characteristics of real life data such as PCR errors and variable amplicon abundances. Table 1 shows a comparison of some of these simulation tools.

**Table 1.** Comparison of metagenomic simulator features. Software management systems cited are apt (24), conda (25), CPAN (26), docker (27) and pip (28). Programs with no management system listed are nevertheless open source and available for download and manual installation and use. Sequencing technologies cited are Illumina, Ion Torrent, Oxford Nanopore Technologies, Pacific Biosciences, Roche 454 (all described by (29)), and Sanger sequencing and ABI SOLiD (30). Illumina layout indicates single ended (SE) or paired end (PE) reads. Output file types are described by (31). Mix. indicates whether the program accepts a list of genomes to simulate a mixed sample. Amps.: can the program simulate amplicon sequencing protocols. VAA: are variable (i.e. non-uniform) amplicon abundances modeled. Ampl. bias: whether differential amplification success is modelled, based on (e.g.) reads’ GC content. Regarding SNP and indel errors, HF denotes high-frequency errors, i.e. recurring position-dependent errors, possibly appearing at the same position across different amplicons, as described in Methods; whereas Reads denotes independent per-read errors, e.g. from sequencing device error. ^1^VLQ is a tool for analyzing wastewater sample sequence data for SARS-CoV-2 variant quantification; it has an unnamed simulation tool available as part of its benchmarking code. ^2^Output from Grinder can be FASTQ-formatted, but the quality scores serve only to indicate whether or not each position incurred a sequencing error.

| Program | Language;<br>Management | Sequencing<br>Technology | Illumina<br>Layout | Output<br>File Type | Mix. | Amps. | VAA | Ampl.<br>Bias | SNP & Indel Errors |  | Quality<br>Scores |
| --- | --- | --- | --- | --- | --- | --- | --- | --- | --- | --- | --- |
|  |  |  |  |  |  |  |  |  | HF | Reads |  |
| SWAMPy | python;<br>docker | Illumina | PE | FASTQ | ✓ | ✓ | ✓ | ✓ | ✓ | ✓ | ✓ |
| ART (14) | C++, Perl;<br>conda | Illumina, 454,<br>SOLiD | PE | FASTQ,<br>SAM | — | ✓ | — | — | — | ✓ | ✓ |
| InSilicoSeq (13) | python;<br>conda, docker, pip | Illumina | PE | FASTQ | ✓ | — | — | ✓ | — | ✓ | ✓ |
| ww_simulations (18) | python;<br>— | Illumina, ONT | PE | FASTQ | ✓ | ✓ | — | — | — | ✓ | ✓ |
| VLQ<sup>1</sup> (15) | python;<br>conda | Illumina | PE | FASTQ | ✓ | — | — | — | — | ✓ | ✓ |
| pIRS (19) | C++, Perl;<br>— | Illumina | PE | FASTQ | — | — | — | ✓ | — | ✓ | ✓ |
| GemSIM (20) | python;<br>— | Illumina, 454 | SE, PE | FASTQ,<br>SAM | ✓ | — | — | — | — | ✓ | ✓ |
| Grinder (7) | Perl;<br>apt, CPAN | Illumina, 454,<br>Sanger | PE | FASTA,<br>FASTQ | ✓ | ✓ | ✓ | — | — | ✓ | —<sup>2</sup> |
| NeSSM (21) | C, Perl;<br>— | Illumina, 454 | SE, PE | FASTQ | ✓ | — | — | ✓ | — | ✓ | ✓ |
| BEAR (22) | Perl, python;<br>— | Illumina, 454,<br>Ion Torrent | SE, PE | FASTQ | ✓ | — | — | — | — | ✓ | ✓ |
| FASTQSim (23) | python;<br>— | Illumina, PacBio,<br>IonTorrent, 454 | SE | FASTQ | ✓ | — | — | — | — | ✓ | ✓ |

Our simulator, SWAMPy (**S**imulating SARS-CoV-2 **W**astewater **A**mplicon **M**etagenomes with **Py**thon), is intended to produce a realistic set of reads that might be generated through multiplex PCR of a wastewater sample, and sequenced by an Illumina sequencer. We model the following scenario:

1. Different viral genomes coming from a human population contaminate wastewater systems, creating a mixture of virus variants which is then captured in a wastewater sample. At this stage, viral RNAs are exposed to RNA degradation and there is a variation in variant abundance in the mixture.
2. After sample collection, PCR amplification of whole viral genomes in segments using a pre-defined primer set results in an amplicon population. At this stage, amplicons gain PCR errors and there is further variation in amplicon abundance due to differential amplification success of the primers of a given primer-set.
3. These amplicons are then sequenced on an Illumina device, creating paired-end reads of a fixed length. At this stage, sequencing errors appear.

## Methods

The overall workflow of SWAMPy can be seen in Fig. 1. The four basic steps of our software pipeline are detailed as follows:

**Fig. 1.**
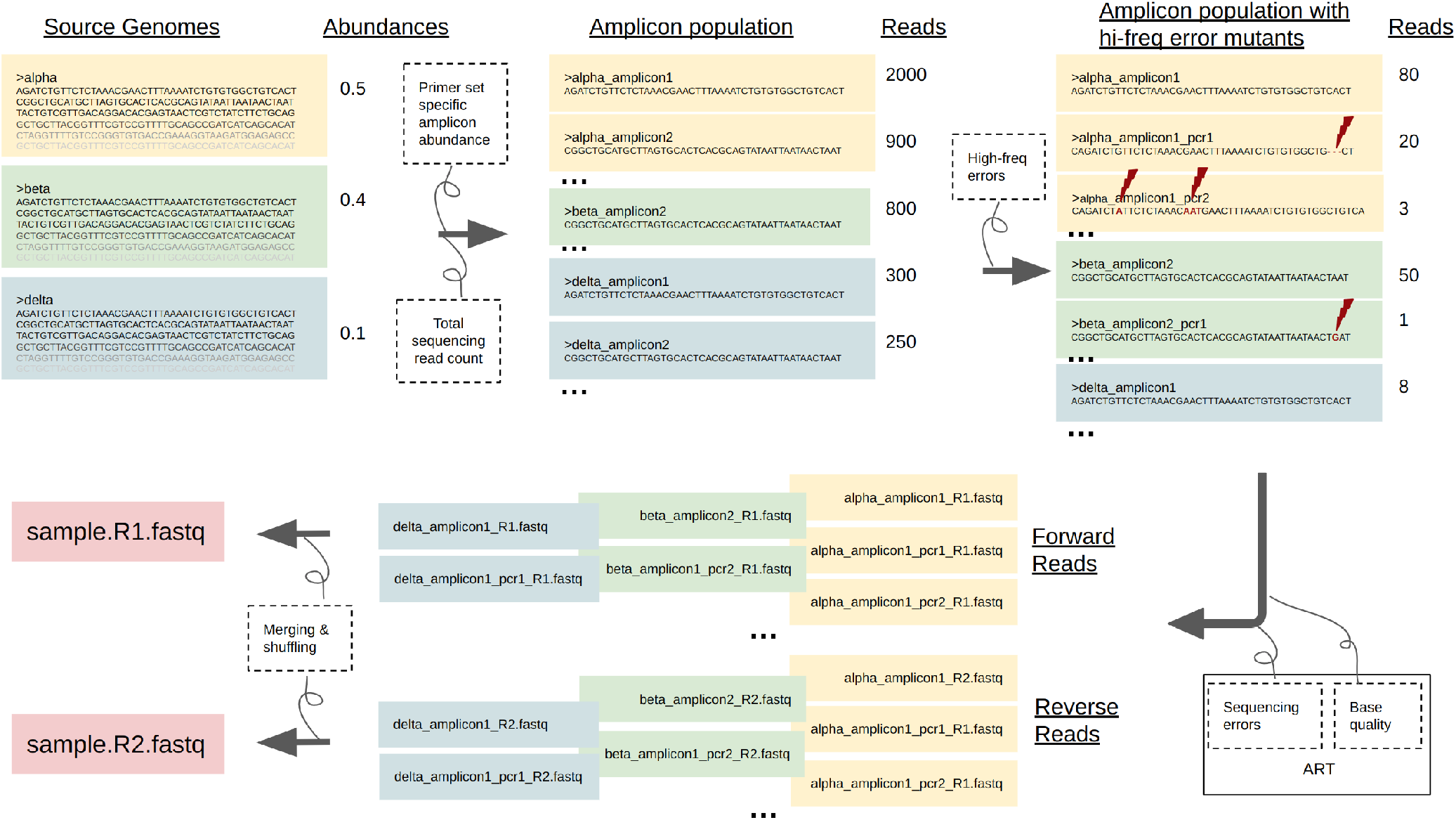
Summary of the SWAMPy workflow. Clockwise from top left, SWAMPy takes as input the genomes of the variants to be represented in the simulated wastewater sample, as well as information on the relative abundances of the variants in the simulated mixture. Source or input genomes are sliced according to a primer set to create a reference amplicon population, and amplicon read depths are adjusted to fit the amplicon abundance distribution of the given primer set (see supplementary material) while taking into account a user-defined parameter which reflects the quality of the samples. The amplicon population is then further diversified by the addition of PCR mutants bearing different kinds of high-frequency errors, using parameters estimated from real data (see supplementary material). The resulting reference and mutant amplicons, with corresponding read counts, are passed to the art_illumina program of ART (14) to model the Illumina sequencing step, where sequencing errors and base qualities are simulated. Finally, reads are merged and shuffled to create mixed-variant forward and reverse FASTQ files.

1. Create an initial amplicon population
2. Simulate the number of DNA fragments (copies) per amplicon
3. Simulate high-frequency errors by mutating amplicons in the amplicon population
4. Simulate sequencing reads using ART

### A. Create an initial amplicon population

The software assumes that the user has supplied a set of SARS-CoV-2 genomes, which we refer to as source genomes, and has selected a primer set among the supported primer sets in the program. The default primer set is ARTIC version 1. On the basis of this selection, amplicons are extracted from each genome as follows. First, we use Bowtie 2 (32) to align the primers (forward and reverse complement) to each virus genome to detect primer binding positions on the source genomes. Next, we slice the source genomes from the primer binding positions to obtain individual amplicons of each source genome, including the primer sequences. During the alignment step, some primers may not align well with the viral genome, and in those cases, the corresponding amplicon is not produced. While this is a strict penalty for primer misalignment, it approximates the observation that some amplicons can be dropped due to mutations in the target regions (33).

### B. Simulate numbers of copies per amplicon

To simulate numbers of copies per amplicon and genome, we offer two versions of a combined Multinomial and Dirichlet model. For either of these models, the user must supply three parameters: total target number of reads *N*, a vector of genome abundances *p*_*g*_indexed over genomes *g*, and a Dirichlet parameter *c*. The choice of *c* roughly equates to a measure of sample quality: a higher value of *c* (e.g. 200) corresponds to high quality samples (roughly uniform abundances of amplicons between simulation runs) and a lower value (e.g. 10) to low quality samples (highly variable amplicon abundances between simulations, with higher rates of amplicon dropout). For the supported primer sets SWAMPy provides an experimentally derived prior on the amplicon proportions *π*_*a*_indexed over amplicons *a*.

**Model 1** (equal expected amplicon proportions across genomes):

1. Sample genome read counts *N*_*g*_ from Multinomial(*N, p*_*g*_)
2. Sample amplicon proportions *p*_*a*_ from Dirichlet(*c* × *π*_*a*_) to be shared across all genomes
3. For each genome, sample numbers of reads per amplicon per genome as *x*_*a,g*_∼ Multinomial(*N*_*g*_, *p*_*a*_)

**Model 2** (different amplicon proportions across genomes):

1. Sample genome read counts *N*_*g*_ from Multinomial(*N, p*_*g*_)
2. For each genome, independently sample amplicon proportions *p*_*a,g*_ from Dirichlet(*c* × *π*_*a*_)
3. For each genome, sample numbers of reads per amplicon per genome as *x*_*a,g*_∼ Multinomial(*N*_*g*_, *p*_*a,g*_)

One subtlety in this process is that the numbers of reads do not account for amplicons dropped in the alignment step, which leads to some missing reads. For example, if the model assigns 100 reads to amplicon 1 in genome A, yet a mutation at the primer site of this amplicon causes it to drop out, then the total number of reads produced would be 100 fewer than expected. Hence the actual total number of reads may be less than the target, *N*.

### C. Add high-frequency errors

While we model sequencing error with standard bioinformatics tools, we developed a new model for the effects of RNA degradation, PCR mutations, and other library preparation artefacts, which we collectively refer to as high-frequency errors. We define high-frequency errors as non-naturally occurring insertions, deletions and substitutions that non-independently affect multiple reads. We further classify high-frequency errors as unique or recurrent with respect to their appearance across different source genomes in the mixture. Recurrent errors are the ones that are present in all source genomes in the simulated mixture, consistent with our observation of individual errors affecting multiple real wastewater sequencing experiments.

These might originate for example from genomic positions particularly susceptible to degradation, or context-dependent PCR errors (34, 35). In contrast, unique high-frequency errors are present in only one of the genomes in the mixture, corresponding for example to PCR errors that are not context-dependent, and low-rate RNA degradation errors.

#### C.1. Sampling high-frequency errors

To simulate high-frequency errors in SWAMPy, we first create a table like that shown in Table 2 containing all the sampled high-frequency errors to be introduced.

**Table 2.**
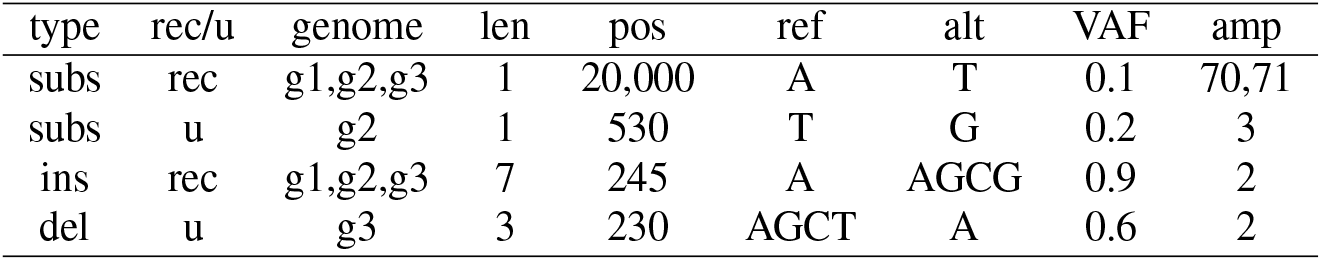
Example simulated high-frequency errors. Abbreviations: rec: recurrent, u: unique, subs: substitution, del: deletion, ins: insertion, amp: amplicon number, len: length, alt: alternative allele, pos: genomic position, g*N* : SARS-CoV-2 variant genome.

1. The number of each type of error to be introduced is sampled from Poisson(*L*× *R*) where *L* = 29903 is the length of the Wuhan reference genome Wuhan-Hu-1 (36) and *R* is the error rate of the given type of error (insertion, deletion, or substitution, each either unique or recurrent). This Poisson distribution approximates the Binomial(*L, R*) distribution since error rates are typically low. Error rates are user-definable for each of the six types of error, with default values estimated from real wastewater experiments (see supplementary material).
2. A genomic position for each error is sampled randomly without replacement from Wuhan-Hu-1. For unique errors, one of the source genomes is randomly assigned with sampling weights equal to the genome abundances in the mixture. Moreover, if more than one amplicon spans the previously determined error position, a unique error is assigned only one of them. Recurrent errors are assigned to all source genomes and overlapping amplicons.
3. An error length is assigned to each error. The error length is always 1 for substitutions, while for deletions it is sampled from a geometric distribution with parameter *n*; higher *n* will result in shorter deletions. For insertions, the error length is sampled from Uniform(*m*) where *m* is the maximum insertion length. Error length parameters *n* and *m* can be defined by the user, with their default values obtained from real data (see supplementary material).
4. An alternative allele is created for each error. For substitutions, it is a random single nucleotide that is different from the reference genome, and for insertions, it is a sequence of randomly sampled nucleotides of the previously determined error length.
5. A variant allele frequency, *f*, is sampled for each error from a Beta(*α, β*) distribution. The Beta distribution parameters are similarly user-definable, separately for unique and recurrent substitutions, insertions and deletions. Assigned VAF values are the expected VAF of the recurrent errors in the final mixture, while for unique errors, the expected value of the VAF in the final mixture will be the product of the assigned VAF *f* and the corresponding amplicon abundance *π*_*a*_.

#### C.2. Apply sampled errors to simulated amplicons

After we compile the table that contains all simulated errors, we process each source genome *g* and each amplicon *a* in the amplicon population that we previously created. For each *a, g*:

1. Errors that affect genome *g* and amplicon *a* are selected from the error table.
2. Because simulated error positions are based on the Wuhan-Hu-1 reference and a variant amplicon in a wastewater sample may contain indels, the amplicon sequences are aligned to Wuhan-Hu-1 using Bowtie 2 (32) and the positions of the errors within the amplicon are determined.
3. For each error *e*, the number of reads in which *e* is present is determined by sampling a read count *n*_*e*_ from Binomial(*x*_*a,g*_, *f*_*e*_), where *x*_*a,g*_ is the total read count of amplicon *a* for genome *g* as described in Section B, and *f*_*e*_ is the VAF of the error as determined in Section C.1.
4. For each possible combination *i* of high-frequency errors affecting genome *g* and amplicon *a*, a read count *n*_*i*_is randomly sampled respecting individual read counts of the errors. We make no attempt to simulate correlations among the errors on amplicons as simulating error inheritance for each amplicon is computationally too expensive and we assume errors on an amplicon are independent.
5. Finally, for each combination *i* of errors affecting *a* and *g*, a new corresponding modified amplicon sequence is created.

### D. Simulate read sequencing using ART

To create a set of simulated paired-end Illumina reads from each amplicon, each with a given read count, we use the program ART (14). We use ART’s paired-end amplicon mode, as well as the noALN and maskN flags. These settings create 150bp paired-end reads, and faithfully transcribe any “N” characters appearing within the amplicons. We use a set of default error rates and quality score profiles tuned for the Illumina MiSeq V3 sequencer, though the ART package has options available for other platforms and read lengths. A full list of the flags used is in the supplementary material.

Finally, we use a custom script based on ubiquitous bash utilities to concatenate all of the FASTQ read files, and shuffle their order to avoid potential biases in case any downstream application software can be influenced by read ordering.

## Results

The source code of our python implementation of SWAMPy, together with the program documentation and exemplar files is available under the GPL-v3 license at https://github.com/goldman-gp-ebi/SWAMPy.

### SWAMPy Implementation

SWAMPy takes as input a multi-FASTA file containing the SARS-CoV-2 variant genomes that will be present in the simulated wastewater sample, as well as a file that contains the relative abundances of these variants in the mixture. For ease of use, other input files (primer-set-specific sequence files, and primer-set-specific amplicon distribution files) were wrapped with a single --primer-set parameter which loads the corresponding input files for the specified primer set. As of December 2022, there are three supported primer sets: ARTIC V1, ARTIC V4 (12) and Nimagen V2 (37). There are many command line parameters that allows fine control of the program such as the parameter *c* that reflects the quality of the wastewater sample as described in Section B, target number of simulated reads, and error rates, VAF and lengths of high-frequency errors as described in Section The full list of command line interface arguments and their explanations are available on the GitHub wiki page: https://github.com/goldman-gp-ebi/SWAMPy/wiki/CLI-arguments.

An example SWAMPy run takes 300 seconds to complete and reaches 700MB of max memory when run with default parameters (three SARS-CoV-2 variants and default error rates and 100,000 total read counts) on a single thread of an Intel Xeon Gold 6336Y 2.40GHz CPU.

SWAMPy produces five output files by default:

- FASTQ files of the simulated forward and reverse reads, matching Illumina standards
- A table that shows the abundance of each wild-type amplicon after the randomness in amplicon copy number sampling (as described in Section B) was applied
- A VCF file that contains all the intended high-frequency errors from the error table described in Section C.1
- A log file

Alignment images of simulation outputs illustrate that the major characteristics of wastewater data are present in the simulated data (Fig. 2) such as overlapping amplicons, different kinds of errors and read depth variation across the genome.

**Fig. 2.**
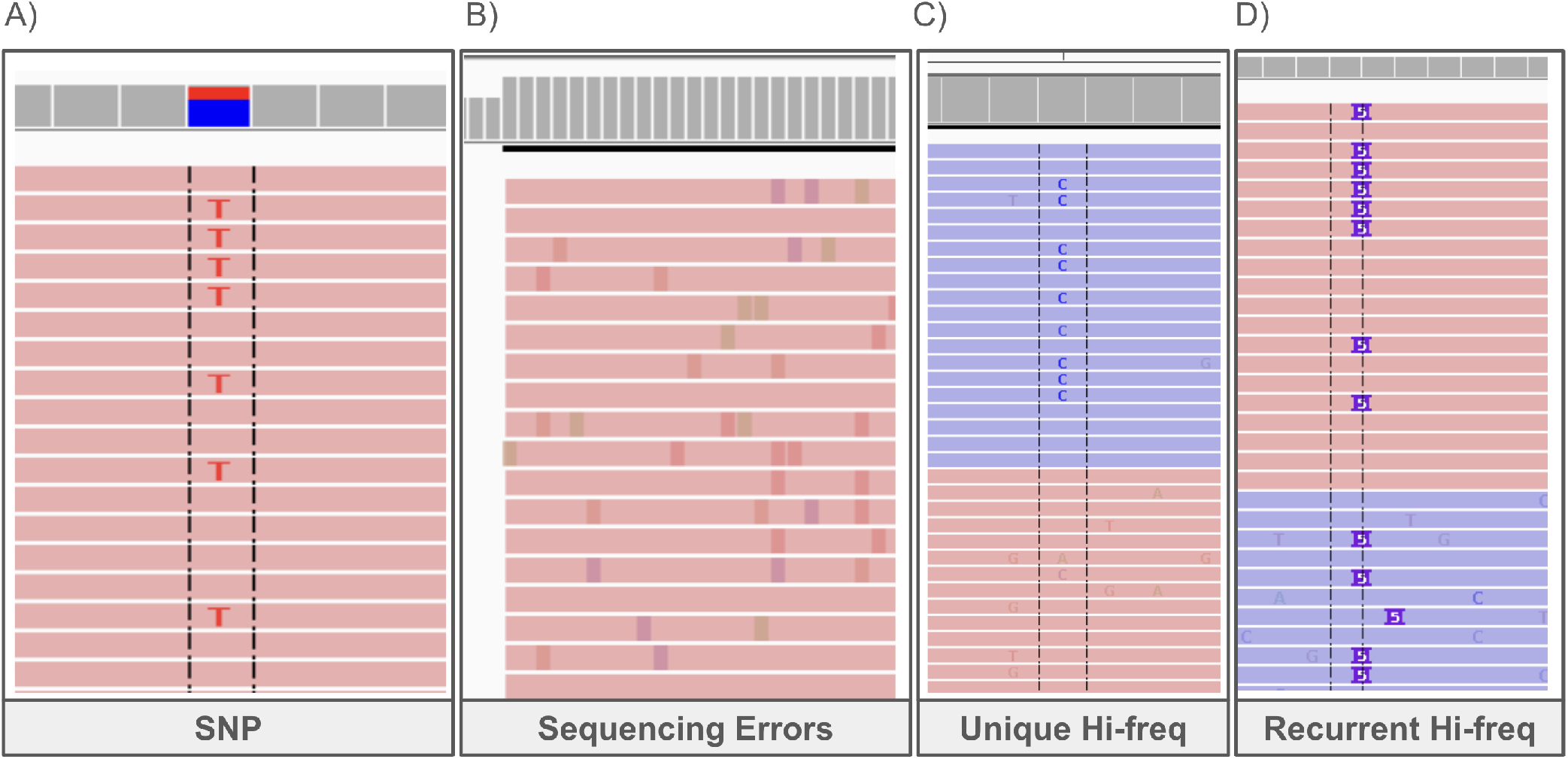
IGV (38) images of SWAMPy simulated reads. Reads come from time point 53 of the simulations described in Results: Use Case, involving SARS-CoV-2 Alpha, Delta and Omicron variants. Uppermost, predominantly grey portion is the coverage track of IGV. Below that, horizontal pink and blue bars show forward and reverse reads, respectively, in “link supplementary alignments” mode of IGV. Black dashed lines are IGV’s optional “center line”, aiding visual perception of alignment of bases; faded colours indicate low quality bases. A) C->T SNP at position 7124 in the Delta variant, not present in the Alpha and Omicron variants. B) Sequencing errors added by ART. This image (zoomed out, with variant bases indicated by colour) is from a read end, where sequencing error density is often higher with Illumina sequencing. Unique high-frequency error. This only appears in one read direction and, despite this exemplar being chosen to be in an amplicon overlap region (not shown), only one of the amplicons carries the error. D) Recurrent high-frequency error (insertion of length 5) appearing in both read directions, and in both amplicons covering the chosen region (not shown).

### Use Case

We used SWAMPy to simulate 73 time points throughout the course of a hypothetical SARS-CoV-2 pandemic where the Alpha (B.1.1.7) variant starts out dominant before Delta (AY.4) rises in frequency and then Omicron (BA.1.1) emerges and takes over (Fig. 3; see the supplementary material for the SWAMPy options used; for the exact abundances at each time point, see supplementary material). Then we used a downstream application program, Freyja, which is designed to detect SARS-CoV-2 variants and their relative abundances from sequencing data obtained from wastewater samples (6, 25, 27, 28). We observe that Freyja is quite successful in demixing the simulated data overall in this relatively complex scenario, though it sometimes inferred the presence of variants that are not specified in the simulated mixture potentially with high frequencies. This probably stems from some high-frequency errors in the simulated data as described in Section C.

**Fig. 3.**
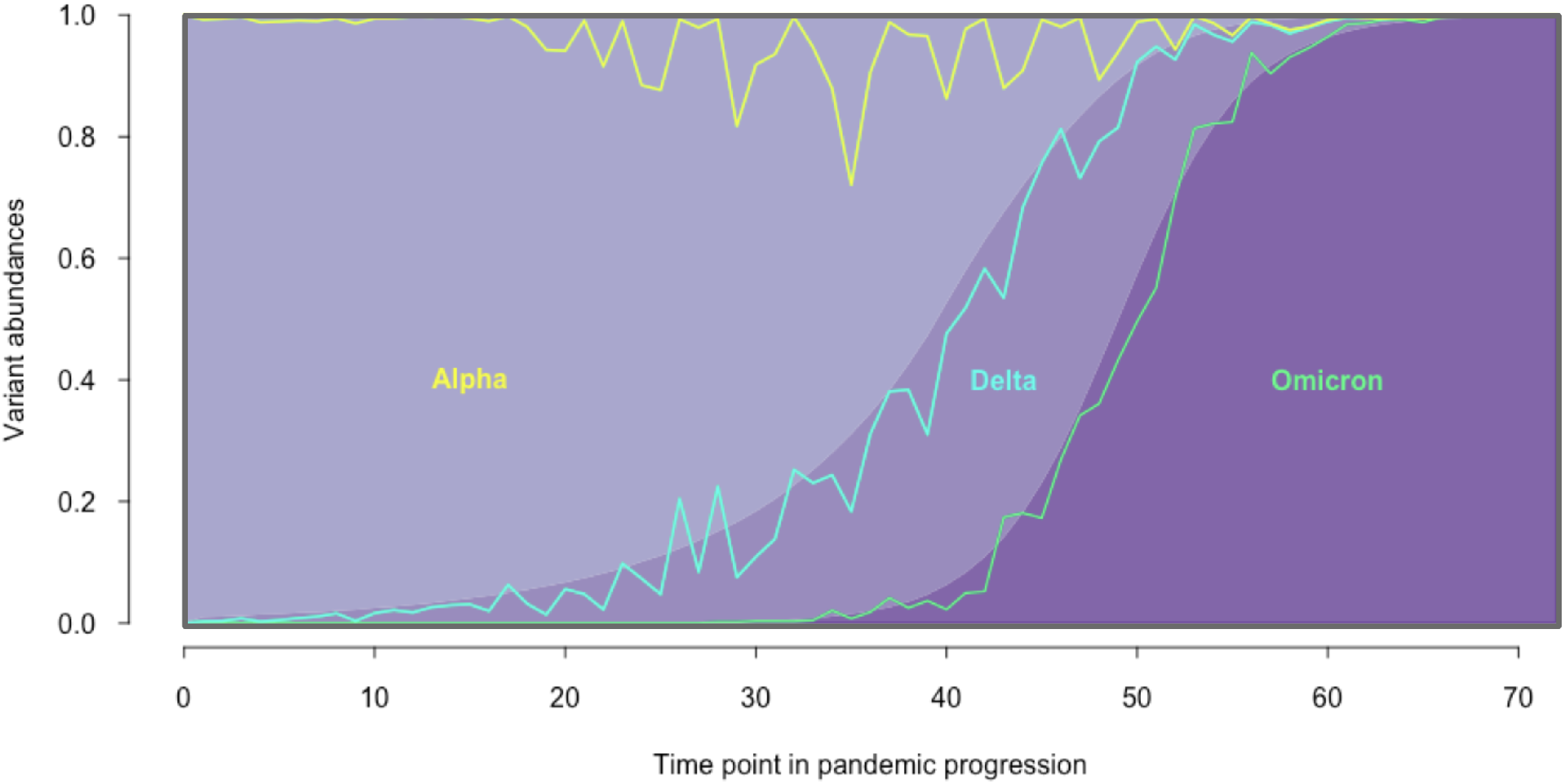
Progression of a simulated pandemic. Sequencing of wastewater samples at 73 time points was simulated with SWAMPy, and corresponding Freyja estimations of SARS-CoV-2 variant abundances were made. Background colors represent the simulated values and lines represent the Freyja estimations. Lines generally follow the boundaries between the shaded areas, suggesting broadly accurate variant proportion estimates from Freyja. The region above the yellow line shows the sum of non-simulated variants (i.e. false positive variant detection) that Freyja erroneously inferred.

## Conclusions

We have shown that SWAMPy is a viable simulation tool for SARS-CoV-2 wastewater metagenomes, building on the simulator ART but much better suited to the modelling challenges idiosyncratic to SARS-CoV-2 metagenomes such as high-frequency errors and irregular amplicon abundance profiles. Both of these models are based on real abundance and error data, from a large number of *in vitro* whole-genome amplicon sequencing experiments, detailed in the supplementary material. Compared with other metagenomic simulators, many of which support a range of complex features such as chimeric amplicons and shotgun metagenomic reads, SWAMPy aims to fill a niche created by metagenomic amplicon studies such as wastewater surveillance of SARS-CoV-2. This niche seems important given the disparity between features available in most general-purpose metagenomic simulators (Table 1) and the requirements of tools being developed for wastewater studies.

Our simulator supports two versions of the ARTIC protocol, which at present is the most prevalent sequencing protocol for SARS-CoV-2 metagenomes, and the Nimagen V2 protocol. We will strive to support future iterations of these, as well as new superseding protocols as they arise in the future. There are other areas where we hope to make improvements to modelling and usability, such as supporting a greater range of sequencing platforms, ensuring that amplicon dropout rates match closely with experimental findings and accounting for some additional parameters in high-frequency error models, which requires a deeper understanding of error mechanisms through controlled experiments.

We hope that this simulation tool will prove valuable by providing non-trivial test cases especially for strain-resolving SARS-CoV-2 metagenomics algorithms, and for creating control case data for researchers working on SARS-CoV-2 wastewater studies. Wastewater surveillance can provide a cheaper alternative to widespread sequencing of clinical SARS-CoV-2 samples, and it is our hope that through appropriate modelling and simulation of the processes involved in amplicon sequencing of wastewater, these data can be leveraged to their full potential in aiding public health.

## ACKNOWLEDGEMENTS

The authors thank Joshua Quick; members of the JBC-led Wastewater Genomics collaboration, in particular Terry Burke, Steve Paterson, Christopher Quince and Sébastien Raguideau; SWAMPy program testers Charlotte West, Lukas Weilguny, Sevim Seda Çokoğ lu, Kivilcim Başak Vural, Melih Yildiz and Gözde Atağ ; and Mehmet Somel who supported this project.

## Supplementary Note 1: Parameter Estimation and Simulation Experiment Parameters

### A. Amplicon abundance estimation

To estimate Dirichlet parameters for amplicon abundances of the AR-TIC v3 and Nimagen v2 primer schemes, we summed amplicon counts over a number of experiments, using results from both synthetic SARS-CoV-2 sequences and real wastewater data. In either case, viral RNA was reverse transcribed before being amplified using each of these two primer schemes. The spreadsheets at https://github.com/goldman-gp-ebi/SWAMPy/tree/main/supplementary_files provides a summary of amplicon counts through a number of experiments. We excluded amplicon counts in experiments where there was an obvious systematic bias: in some experiments one of the two primer pools failed, and we discarded these results. Other amplicons failed to amplify in the synthetic virus genomes; this was due to the synthetic genomes being produced in 5 kbp chunks. We again discarded these amplicon counts. Finally we normalised our counts’ values so that they summed to 1.

For the Artic v4 primer scheme, we used an amplicon abundance profile based on a single experiment (Joshua Quick, personal communication). The coverage file for this experiment is again provided at https://github.com/goldman-gp-ebi/SWAMPy/tree/main/supplementary_files.

### B. ART Parameters

To simulate sequencing errors we used the program **art_illumina** (part of the ART suite of simulation tools (14)), with the following command-line parameters:

~~~
-amplicon
-paired
-noALN
-maskN 0
-seqSys SEQ_SYS
-len READ_LENGTH
-rcount NUMBER_OF_READS
-rndSeed SEED
-in INPUT_AMPLICON_FASTA
-out OUTPUT_FASTQ_FILENAME
~~~

The variables SEQ_SYS, READ_LENGTH, and NUMBER_OF_READS are set by the user. SEED, INPUT_AMPLICON_FASTA, and OUTPUT_FASTQ_FILENAME are changed for each amplicon of each genome that is being simulated.

To define the default high-frequency error rates in SWAMPy, we performed an error characterisation analysis on 121 real wastewater sequencing datasets. Wastewater sequencing experiments were conducted by members of JBC-led Wastewater Genomics collaboration. 12 of the 121 samples are mixtures of synthetically produced SARS-CoV-2 variant genomes, which mimic wastewater samples. Their ENA accessions are: ERR10084556, ERR10084565, ERR10084558, ERR10084543, ERR10084590, ERR10084581, ERR10084585, ERR10084577, ERR10084594, ERR10084592, ERR10084549, ERR10084547, ERR10084599, ERR10084595, ERR10084586, ERR10084579, ERR10084589, ERR10084564, ERR10084588, ERR10084562. 109 of them were sampled from wastewater in the UK and the data will be publicly available soon.

For each sample of these datasets, we mapped raw reads to the Wuhan-Hu-1 (36) reference genome using Bowtie 2.4.4 (32). We then used bcftools mpileup 1.13 (39) to obtain VCF files. We did not perform a separate variant calling as we are interested in errors as opposed to SNPs and needed every discrepancy between the reference genome and our sample reads; henceforth will refer to such differences as “variants”. We filtered out positions with a read depth (DP) *<* 10 and with *<* 5 reads supporting the alternative allele (AD). The remaining variants are classified into different categories as summarised in Fig. 4. First, if the variant allele frequency (VAF) of a variant is low, in particular lower than 0.02, we filtered out that variant since such low-frequency variants might be the result of standard sequencing errors (40) and are unlikely to affect downstream analyses. The remaining variants included real polymorphisms between different SARS-CoV-2 variants, as well as possible high-frequency errors, which is the group of errors of interest. To identify likely high-frequency errors, we focused on putative nonviable mutations, since nonviable mutations cannot be real polymorphisms.

**Fig. 4.**
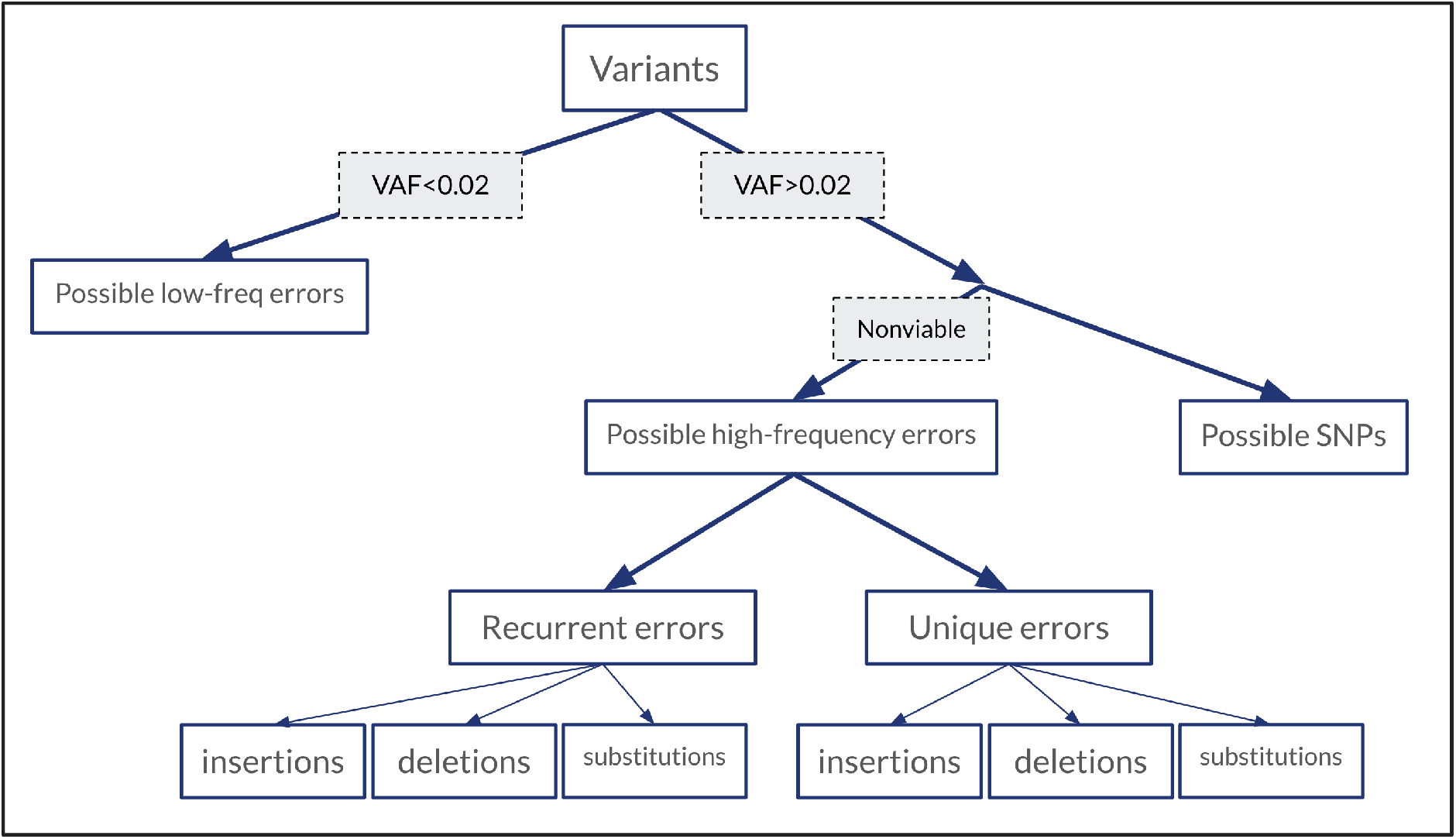
Variant classification criteria.

### C. High-frequency errors parameter estimation

We define putative nonviable mutations as nonsense (stop codon) substitutions or indels of length not a multiple of three found on ORF1ab and S open reading frames. We excluded other smaller and less characterized genes from the analysis since these can present viable nonsense mutations (41, 42). To further eliminate possible real polymorphisms, we also excluded a portion from the 3^′^ends of the ORF1ab and S open reading frames since nonsense mutations could be tolerable there. The exact positions included are 266–12000, 13465–20000 and 21563–25000. We further divided the high-frequency errors as either recurrent or unique based on if they appeared in more than one wastewater sample.

We subdivided both recurrent and unique errors into insertions, deletions and substitutions subclasses, and estimated default rates of these events by counting the observed variants in each class and normalizing by the number of genome positions with sufficient coverage and correcting for the expected proportions of high-frequency errors that would not have been identified (indels with length multiple of three and non-stop codon substitutions). A more exhaustive description of the approach to infer high-frequency error rates and lengths is given in the following subsections.

#### C.1. Indel error length distributions

Following the length distributions observed in our putative high-frequency indel errors (see figure 5), we model high-frequency indel error lengths as:

**Fig. 5.**
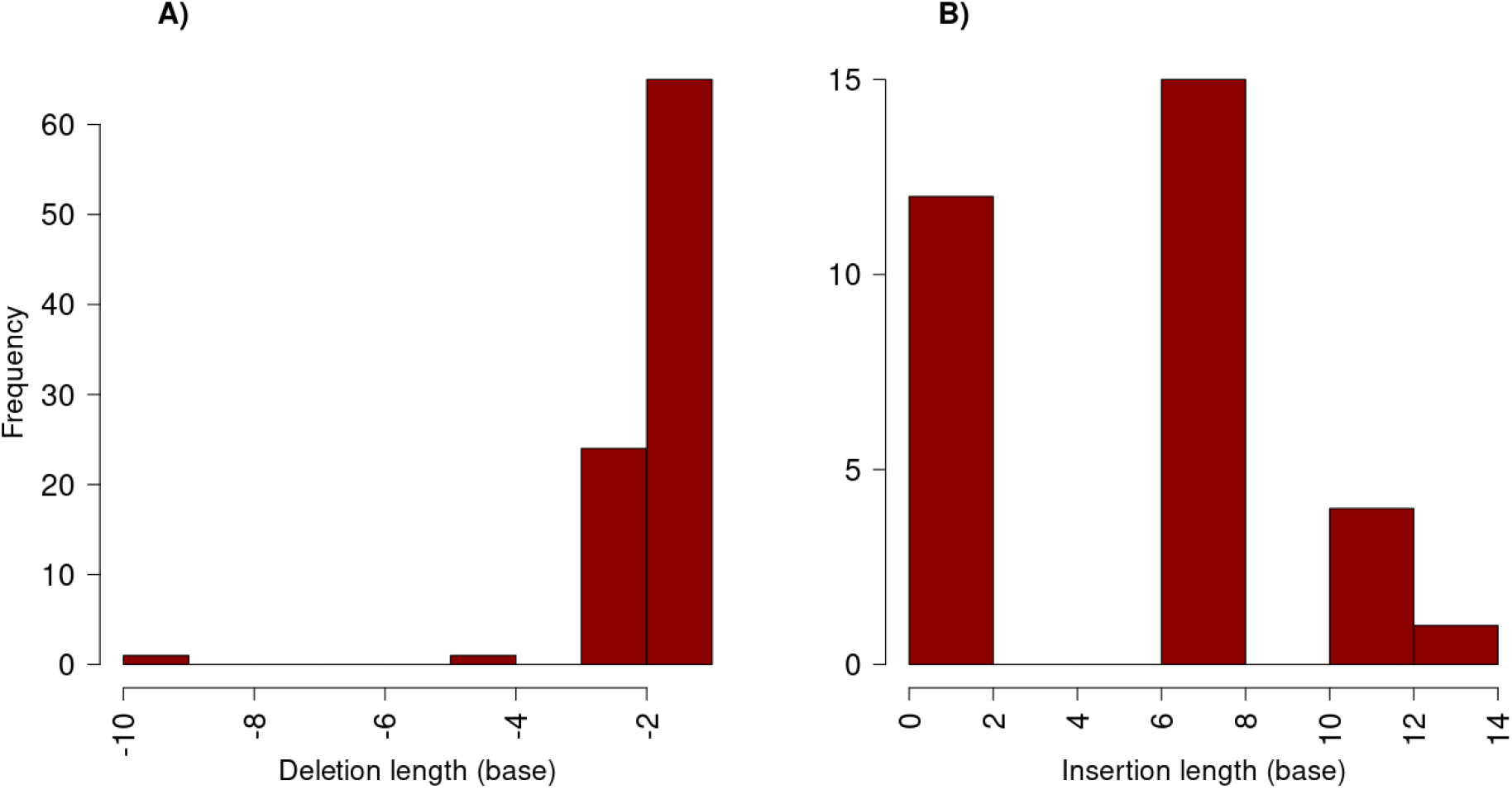
Histograms of putative indel error lengths observed in real data. A) Deletions; B) Insertions.

- Insertions: uniform distribution with minimum length 1 and maximum 14 (the minimum and maximum lengths found in real data) *U* (1, 14).
- Deletions: geometric distribution with parameter *p*, Geometric(*p*).

We could not use a standard distribution fitting technique to estimate the indel length geometric distribution parameter *p* because we cannot observe putative high-frequency error deletions with length multiple of three. Instead, to avoid potential biases due to the missing data, we estimate *p* using only the counts of putative high-frequency deletion errors of length one and two. Since under Geometric(*p*) the ratio of the probability of lengths *l* = 2 and *l* = 1 is

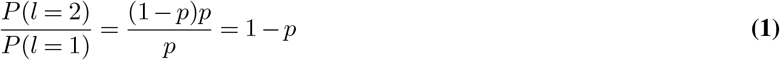

We use the estimator

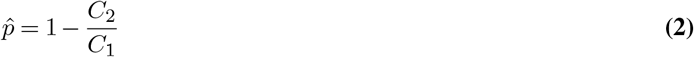

where *C*_2_ and *C*_1_ are the observed counts of putative high-frequency deletion errors of length 2 and 1 respectively.

#### C.2. High-frequency error rate estimation

We estimated six separate high-frequency error rates, one for each of the six classes of high-frequency errors. For each high-frequency error class *i*, we call *C*_*i*_ its putative error count observed in the real data, and we estimate its rate *r*_*i*_ as

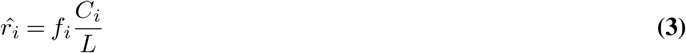

where *f*_*i*_ is the missing data correction factor for class *i*, and *L* is the total number of genome positions, across all real datasets, that we considered when looking for putative high-frequency errors. For substitutions, *f*_*i*_ corrects for the fact that only nonsense variants could be identified as putative high-frequency errors, while high-frequency errors causing other types of substitutions were not counted as they could not be distinguished from real polymorphisms. Consequently, we defined the correction factor *f*_*i*_ for substitutions as *M/M*_*n*_, where *M* is the number of all possible substitutions across the considered portion of the reference SARS-CoV-2 genome (Wuhan-Hu-1, 36), so *M* = 3*L* where *L* is the length of the considered portion of reference genome; and *M*_*n*_ is the number of possible nonsense mutations across the same portion of the reference genome.

For insertions, the correction factor *f*_*i*_ is simply 3*/*2 because we assume a uniform high-frequency insertion length distribution and because we did not count insertion with length multiple of three among putative high-frequency insertion errors.

For the deletion correction factor we took a similar approach, but accounting for the assumed geometric distribution of lengths. In a geometric distribution with parameter *p*, the total probability of all multiples of three is given by

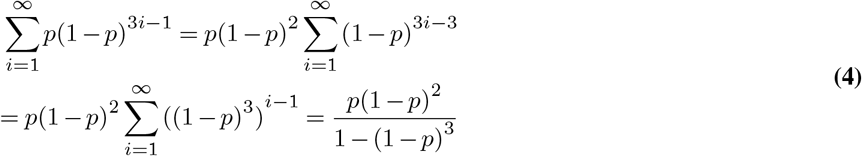

where for the last step we used the identity 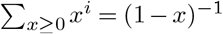. The total probability of the lengths that we do observe is 1 minus this value, i.e. 1 − *p*(1 − *p*)2*/*(1 − (1 − *p*)^3^), and therefore we use as correction factor *f*_*i*_ of high-frequency deletions its inverse:

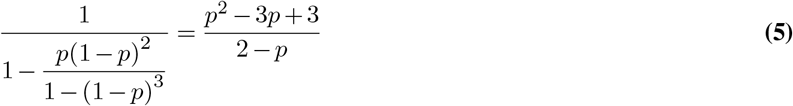

#### C.3. High-frequency error frequency distributions

To model the default variant allele frequencies (VAF) of high-frequency errors of each type, we use the Beta distribution (see e.g. 43) whose default parameters were estimated from the frequencies of putative high-frequency errors using the method of moments.

### D. Use-case

The exact proportions of the SARS-CoV-2 variants at the 73 time points in our simulations are shown in Table 3.

**Table 3.**
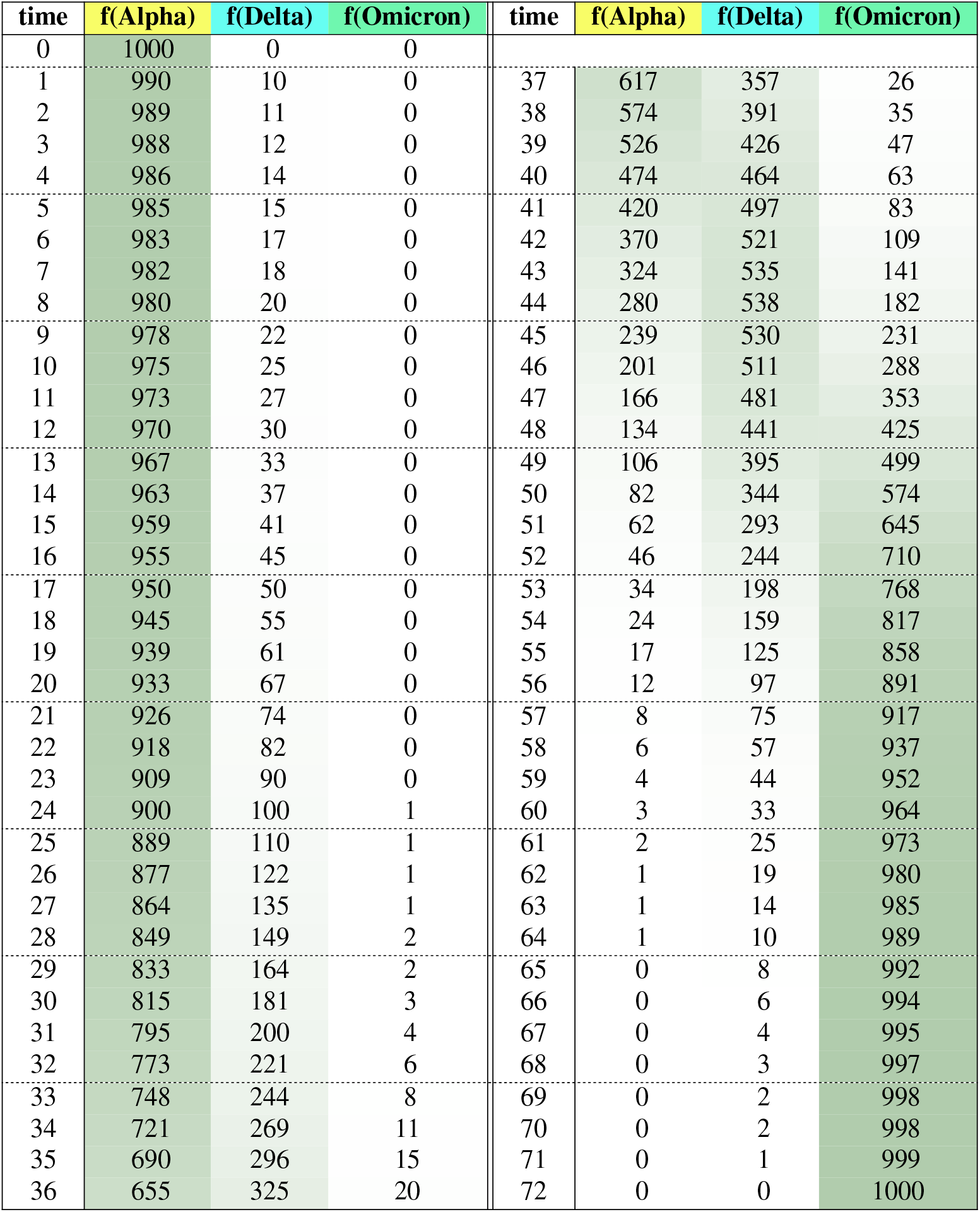
Simulated genome proportions.

The non-default SWAMPy options used to simulate these 73 time points are:

~~~
−ins 0.0002
−del 0.00115
−subs 0.005
−rins 0.0002
−rdel 0.00115
−subs 0.005
−amplicon_distribution dirichlet_2
−amplicon_pseudocounts 200
~~~

## Notes

https://github.com/goldman-gp-ebi/SWAMPy

### Competing Interest Statement

The authors have declared no competing interest.

### Summary of Updates

Minor clarifications and correction of typos. Addition of 1 table, comparing features of available metagenomics simulators with those of our program SWAMPy. Addition of H. Denise as co-author to correctly reflect contribution to this project.

## Bibliography

1. Mathew R. Brown, Matthew J. Wade, Shannon McIntyre-Nolan, Irene Bassano, Hubert Denise, David Bass, John Bentley, Joshua T. Bunce, Jasmine Grimsley, Alwyn Hart, Till Hoffmann, Aaron Jeffries, Steve Paterson, Mark Pollock, Jonathan Porter, David Smith, Ronny van Aerle, Glenn Watts, Andrew Engeli, and Gideon Henderson. Wastewater Monitoring of SARS-CoV-2 Variants in England: Demonstration Case Study for Bristol (Dec 2020 – March 2021). ePrints, Apr. 2021. doi: 10.13140/RG.2.2.27215.00162.

2. Véronique Hourdel, Aurelia Kwasiborski, Charlotte Balière, Séverine Matheus, Christophe Frédéric Batéjat, Jean-Claude Manuguerra, Jessica Vanhomwegen, and Valérie Caro. Rapid genomic characterization of SARS-CoV-2 by direct amplicon-based sequencing through comparison of MinION and Illumina iSeq100™ system. Frontiers in Microbiology, 11:571328, Sep. 2020. doi: 10.3389/fmicb.2020.571328.

3. Devon A. Gregory, Chris G. Wieberg, Jeff Wenzel, Chung Ho Lin, and Marc C. Johnson. Monitoring SARS-CoV-2 populations in wastewater by amplicon sequencing and using the novel program SAM Refiner. Viruses, 13:1647, Aug. 2021. doi: 10.3390/V13081647.

4. Katharina Jahn, David Dreifuss, Ivan Topolsky, Anina Kull, Pravin Ganesanandamoorthy, Xavier Fernandez-Cassi, Carola Bänziger, Alexander J. Devaux, Elyse Stachler, Lea Caduff, Federica Cariti, Alex Tuñas Corzón, Lara Fuhrmann, Chaoran Chen, Kim Philipp Jablonski, Sarah Nadeau, Mirjam Feldkamp, Christian Beisel, Catharine Aquino, Tanja Stadler, Christoph Ort, Tamar Kohn, Timothy R. Julian, and Niko Beerenwinkel. Early detection and surveillance of SARS-CoV-2 genomic variants in wastewater using COJAC. Nature Microbiology, 7:1151–1160, Jul. 2022. doi: 10.1038/s41564-022-01185-x.

5. Renan Valieris, Rodrigo D Drummond, Alexandre Defelicibus, Emmanuel Dias-Neto, Rafael A Rosales, and Israel Tojal da Silva. A mixture model for determining SARS-CoV-2 variant composition in pooled samples. Bioinformatics, 38:1809–1815, Feb. 2022. doi: 10.1093/bioinformatics/btac047.

6. Smruthi Karthikeyan, Joshua I. Levy, Peter De Hoff, Greg Humphrey, Amanda Birmingham, Kristen Jepsen, Sawyer Farmer, Helena M. Tubb, Tommy Valles, Caitlin E. Tribelhorn, Rebecca Tsai, Stefan Aigner, Shashank Sathe, Niema Moshiri, Benjamin Henson, Adam M. Mark, Abbas Hakim, Nathan A. Baer, Tom Barber, Pedro Belda-Ferre, Marisol Chacón, Willi Cheung, Evelyn S. Cresini, Emily R. Eisner, Alma L. Lastrella, Elijah S. Lawrence, Clarisse A. Marotz, Toan T. Ngo, Tyler Ostrander, Ashley Plascencia, Rodolfo A. Salido, Phoebe Seaver, Elizabeth W. Smoot, Daniel McDonald, Robert M. Neuhard, Angela L. Scioscia, Alysson M. Satterlund, Elizabeth H. Simmons, Dismas B. Abelman, David Brenner, Judith C. Bruner, Anne Buckley, Michael Ellison, Jeffrey Gattas, Steven L. Gonias, Matt Hale, Faith Hawkins, Lydia Ikeda, Hemlata Jhaveri, Ted Johnson, Vince Kellen, Brendan Kremer, Gary Matthews, Ronald W. McLawhon, Pierre Ouillet, Daniel Park, Allorah Pradenas, Sharon Reed, Lindsay Riggs, Alison Sanders, Bradley Sollenberger, Angela Song, Benjamin White, Terri Winbush, Christine M. Aceves, Catelyn Anderson, Karthik Gangavarapu, Emory Hufbauer, Ezra Kurzban, Justin Lee, Nathaniel L. Matteson, Edyth Parker, Sarah A. Perkins, Karthik S. Ramesh, Refugio Robles-Sikisaka, Madison A. Schwab, Emily Spencer, Shirlee Wohl, Laura Nicholson, Ian H. McHardy, David P. Dimmock, Charlotte A. Hobbs, Omid Bakhtar, Aaron Harding, Art Mendoza, Alexandre Bolze, David Becker, Elizabeth T. Cirulli, Magnus Isaksson, Kelly M. Schiabor Barrett, Nicole L. Washington, John D. Malone, Ashleigh Murphy Schafer, Nikos Gurfield, Sarah Stous, Rebecca Fielding-Miller, Richard S. Garfein, Tommi Gaines, Cheryl Anderson, Natasha K. Martin, Robert Schooley, Brett Austin, Duncan R. MacCannell, Stephen F. Kingsmore, William Lee, Seema Shah, Eric McDonald, Alexander T. Yu, Mark Zeller, Kathleen M. Fisch, Christopher Longhurst, Patty Maysent, David Pride, Pradeep K. Khosla, Louise C. Laurent, Gene W. Yeo, Kristian G. Andersen, and Rob Knight. Wastewater sequencing reveals early cryptic SARS-CoV-2 variant transmission. Nature, 609:101–108, Sep. 2022. doi: 10.1038/s41586-022-05049-6.

7. Florent E. Angly, Dana Willner, Forest Rohwer, Philip Hugenholtz, and Gene W. Tyson. Grinder: a versatile amplicon and shotgun sequence simulator. Nucleic Acids Research, 40:e94, Jul. 2012. doi: 10.1093/NAR/GKS251.

8. Merly Escalona, Sara Rocha, and David Posada. A comparison of tools for the simulation of genomic next-generation sequencing data. Nature Reviews Genetics, 17:459–469, Jun. 2016. doi: 10.1038/nrg.2016.57.

9. Yatish Turakhia, Nicola De Maio, Bryan Thornlow, Landen Gozashti, Robert Lanfear, Conor R Walker, Angie S Hinrichs, Jason D Fernandes, Rui Borges, Greg Slodkowicz, Lukas Weilguny, David Haussler, Nick Goldman, and Russell Corbett-Detig. Stability of SARS-CoV-2 phylogenies. PLoS Genetics, 16:e1009175, Nov. 2020. doi: 10.1371/journal.pgen.1009175.

10. Damien Jacot, Trestan Pillonel, Gilbert Greub, and Claire Bertelli. Assessment of SARS-CoV-2 genome sequencing: quality criteria and low-frequency variants. Journal of Clinical Microbiology, 59:e00944–21, 2021. doi: 10.1128/JCM.00944-21.

11. Nicola De Maio, Conor Walker, Rui Borges, Lukas Weilguny, Greg Slodkowicz, and Nick Goldman. Issues with SARS-CoV-2 sequencing data. http://virologi-cal.org; https://virological.org/t/issues-with-sars-cov-2-sequencing-data/473 (accessed 10 Jul. 2023), May 2020.

12. John R Tyson, Phillip James, David Stoddart, Natalie Sparks, Arthur Wickenhagen, Grant Hall, Ji Hyun Choi, Hope Lapointe, Kimia Kamelian, Andrew D Smith, Natalie Prystajecky, Ian Goodfellow, Sam J Wilson, Richard Harrigan, Terrance P Snutch, Nicholas J Loman, and Joshua Quick. Improvements to the ARTIC multiplex PCR method for SARS-CoV-2 genome sequencing using nanopore. bioRxiv, Sep. 2020. doi: 10.1101/2020.09.04.283077.

13. Hadrien Gourlé, Oskar Karlsson-Lindsjö, Juliette Hayer, and Erik Bongcam-Rudloff. Simulating Illumina metagenomic data with InSilicoSeq. Bioinformatics, 35:521–522, Jul. 2018. doi: 10.1093/bioinformatics/bty630.

14. Weichun Huang, Leping Li, Jason R. Myers, and Gabor T. Marth. ART: a next-generation sequencing read simulator. Bioinformatics, 28:593–594, Dec. 2011. doi: 10.1093/bioinformatics/btr708.

15. Jasmijn A. Baaijens, Alessandro Zulli, Isabel M. Ott, Ioanna Nika, Mart J. van der Lugt, Mary E. Petrone, Tara Alpert, Joseph R. Fauver, Chaney C. Kalinich, Chantal B. F. Vogels, Mallery I. Breban, Claire Duvallet, Kyle A. McElroy, Newsha Ghaeli, Maxim Imakaev, Malaika F. Mckenzie-Bennett, Keith Robison, Alex Plocik, Rebecca Schilling, Martha Pierson, Rebecca Littlefield, Michelle L. Spencer, Birgitte B. Simen, Ahmad Altajar, Anderson F. Brito, Anne E. Watkins, Anthony Muyombwe, Caleb Neal, Chen Liu, Christopher Castaldi, Claire Pearson, David R. Peaper, Eva Laszlo, Irina R. Tikhonova, Jafar Razeq, Jessica E. Rothman, Jianhui Wang, Kaya Bilguvar, Linda Niccolai, Madeline S. Wilson, Margaret L. Anderson, Marie L. Landry, Mark D. Adams, Pei Hui, Randy Downing, Rebecca Earnest, Shrikant Mane, Steven Murphy, William P. Hanage, Nathan D. Grubaugh, Jordan Peccia, and Michael Baym. Lineage abundance estimation for SARS-CoV-2 in wastewater using transcriptome quantification techniques. Genome Biology, 23:236, Nov. 2022. doi: 10.1186/s13059-022-02805-9.

16. Askar Gafurov, Andrej Baláž, Fabian Amman, Kristína Boršová Viktória Čabanová, Boris Klempa, Andreas Bergthaler, Tomáš Vinař, and Broňa Brejová. VirPool: model-based estimation of SARS-CoV-2 variant proportions in wastewater samples. BMC Bioinformatics, 23:551, Dec. 2022. doi: 10.1186/s12859-022-05100-3.

17. Nicolae Sapoval, Yunxi Liu, Esther G. Lou, Loren Hopkins, Katherine B. Ensor, Rebecca Schneider, Lauren B. Stadler, and Todd J. Treangen. Enabling accurate and early detection of recently emerged SARS-CoV-2 variants of concern in wastewater. Nature Communications, 14:2834, May 2023. doi: 10.1038/s41467-023-38184-3.

18. Kayikcioglu T, Amirzadegan J, Rand H, Tesfaldet B, Timme RE, and Pettengill JB. Performance of methods for SARS-CoV-2 variant detection and abundance estimation within mixed population samples. PeerJ, 11:e14596, Jan. 2023. doi: 10.7717/peerj.14596.

19. Xuesong Hu, Jianying Yuan, Yujian Shi, Jianliang Lu, Binghang Liu, Zhenyu Li, Yanxiang Chen, Desheng Mu, Hao Zhang, Nan Li, Zhen Yue, Fan Bai, Heng Li, and Wei Fan. pIRS: Profile-based Illumina pair-end reads simulator. Bioinformatics, 28(11):1533–1535, Apr. 2012. doi: 10.1093/bioinformatics/bts187.

20. Kerensa E. McElroy, Fabio Luciani, and Torsten Thomas. GemSIM: general, error-model based simulator of next-generation sequencing data. BMC Genomics, 13(1):74, Feb. 2012. doi: 10.1186/1471-2164-13-74.

21. Jia B, Xuan L, Cai K, Hu Z, Ma L, and Wei C. NeSSM: a next-generation sequencing simulator for metagenomics. PLoS One, 8(10):e75448, Oct. 2013. doi: 10.1371/journal.pone.0075448.

22. Stephen Johnson, Brett Trost, Jeffrey R. Long, Vanessa Pittet, and Anthony Kusalik. A better sequence-read simulator program for metagenomics. BMC Bioinformatics, 15:S14, Sep. 2014. doi: 10.1186/1471-2105-15-S9-S14.

23. Anna Shcherbina. FASTQSim: platform-independent data characterization and in silico read generation for NGS datasets. BMC Research Notes, 7:533, Aug. 2014. doi: 10.1186/1756-0500-7-533.

24. The Debian Project. Apt. https://www.debian.org/doc/manuals/debian-reference/ch02.en.html, 2023. Accessed: 6 Jul. 2023.

25. Anaconda, Inc. Conda. https://docs.conda.io/en/latest/, 2023. Accessed: 29 Jun. 2023.

26. Jarkko Hietaniemi. CPAN. https://www.cpan.org/, 2023. Accessed: 6 Jul. 2023.

27. Docker Inc. Docker. https://www.docker.com, 2023. Accessed: 29 Jun. 2023.

28. The pip developers. Pip. https://pip.pypa.io/en/stable/, 2023. Accessed: 29 Jun. 2023.

29. Barton E. Slatko, Andrew F. Gardner, and Frederick M. Ausubel. Overview of next-generation sequencing technologies. Current Protocols in Molecular Biology, 122:e59, 2018. doi: 10.1002/cpmb.59.

30. Martin Kircher and Janet Kelso. High-throughput DNA sequencing — concepts and limitations. BioEssays, 32:524–536, 2010. doi: 10.1002/bies.200900181.

31. Hongen Zhang. Overview of sequence data formats. In Ewy Mathé and Sean Davis, editors, Statistical Genomics: Methods and Protocols, pages 3–17. Springer New York, New York, NY, 2016. ISBN 978-1-4939-3578-9. doi: 10.1007/978-1-4939-3578-9_1.

32. Ben Langmead and Steven L. Salzberg. Fast gapped-read alignment with Bowtie 2. Nature Methods, 9:357–359, Apr. 2012. doi: 10.1038/nmeth.1923.

33. Aziz Sheikh, Jim McMenamin, Bob Taylor, and Chris Robertson. SARS-CoV-2 Delta VOC in Scotland: demographics, risk of hospital admission, and vaccine effectiveness. The Lancet, 397:2461–2462, Jun. 2021. doi: 10.1016/S0140-6736(21)01358-1.

34. Andreas Meyerhans, Jean Pierre Vartanian, and Simon Wain-Hobson. DNA recombination during PCR. Nucleic Acids Research, 18:1687–1691, Jul. 1990. doi: 10.1093/nar/18.7.1687.

35. Vladimir Potapov and Jennifer L. Ong. Examining sources of error in PCR by single-molecule sequencing. PLoS ONE, 12:e0169774, Jan. 2017. doi: 10.1371/journal.pone.0169774.

36. Fan Wu, Su Zhao, Bin Yu, Yan Mei Chen, Wen Wang, Zhi Gang Song, Yi Hu, Zhao Wu Tao, Jun Hua Tian, Yuan Yuan Pei, Ming Li Yuan, Yu Ling Zhang, Fa Hui Dai, Yi Liu, Qi Min Wang, Jiao Jiao Zheng, Lin Xu, Edward C. Holmes, and Yong Zhen Zhang. A new coronavirus associated with human respiratory disease in China. Nature, 579:265, Mar. 2020. doi: 10.1038/S41586-020-2008-3.

37. Jordy P.M. Coolen, Femke Wolters, Alma Tostmann, Lenneke F.J. van Groningen, Chantal P. Bleeker-Rovers, Edward C.T.H. Tan, Nannet van der Geest-Blankert, Jeannine L.A. Hautvast, Joost Hopman, Heiman F.L. Wertheim, Janette C. Rahamat-Langendoen, Marko Storch, and Willem J.G. Melchers. SARS-CoV-2 whole-genome sequencing using reverse complement PCR: for easy, fast and accurate outbreak and variant analysis. Journal of Clinical Virology, 144:1386–6532, Nov. 2021. doi: 10.1016/j.jcv.2021.104993.

38. James T. Robinson, Helga Thorvaldsdóttir, Wendy Winckler, Mitchell Guttman, Eric S. Lander, Gad Getz, and Jill P. Mesirov. Integrative genomics viewer. Nature Biotechnology, 29: 24–26, Jan. 2011. doi: 10.1038/nbt.1754.

39. Petr Danecek, James K Bonfield, Jennifer Liddle, John Marshall, Valeriu Ohan, Martin O Pollard, Andrew Whitwham, Thomas Keane, Shane A McCarthy, Robert M Davies, and Heng Li. Twelve years of SAMtools and BCFtools. GigaScience, 10, Feb. 2021. doi: 10.1093/gigascience/giab008.

40. Nicholas Stoler and Anton Nekrutenko. Sequencing error profiles of Illumina sequencing instruments. NAR Genomics and Bioinformatics, 3: lqab019, Mar. 2021. doi: 10.1093/nargab/lqab019.

41. Serena Delbue, Sarah D’Alessandro, Lucia Signorini, Maria Dolci, Elena Pariani, Michele Bianchi, Stefania Fattori, Annalisa Modenese, Cristina Galli, Ivano Eberini, and Pasquale Ferrante. Isolation of SARS-CoV-2 strains carrying a nucleotide mutation, leading to a stop codon in the orf 6 protein. Emerging Microbes & Infections, 10:252–255, Feb. 2021. doi: 10.1080/22221751.2021.1884003.

42. Irwin Jungreis, Rachel Sealfon, and Manolis Kellis. SARS-CoV-2 gene content and COVID-19 mutation impact by comparing 44 Sarbecovirus genomes. Nature Communications, 12: 2642, May 2021. doi: 10.1038/s41467-021-22905-7.

43. Kenneth Lange. Applications of the Dirichlet distribution to forensic match probabilities. Genetica, 96:107–117, Jun. 1995. doi: 10.1007/BF01441156.

